# SGI: Automatic clinical subgroup identification in omics datasets

**DOI:** 10.1101/2021.03.12.435108

**Authors:** Mustafa Buyukozkan, Karsten Suhre, Jan Krumsiek

## Abstract

**Summary:** The ‘Subgroup Identification’ (SGI) toolbox provides an algorithm to automatically detect clinical subgroups of samples in large-scale omics datasets. It is based on hierarchical clustering trees in combination with a specifically designed association testing and visualization framework that can process an arbitrary number of clinical parameters and outcomes in a systematic fashion. A multi-block extension allows for the simultaneous use of multiple omics datasets on the same samples. In this paper, we describe the functionality of the toolbox and demonstrate an application example on a blood metabolomics dataset with various clinical biochemistry readouts in a type 2 diabetes case-control study.

**Availability and implementation:** SGI is an open-source package implemented in R. Package source codes and hands-on tutorials are available at https://github.com/krumsieklab/sgi. The QMdiab metabolomics data is included in the package and can be downloaded from https://doi.org/10.6084/m9.figshare.5904022.

## 1 Introduction

The identification of patient subgroups from high-dimensional molecular profiles has become a central approach in biomedical research, driven by the wide availability of modern “multi-omics” datasets [1–3]. The central idea is that genomics, transcriptomics, proteomics, metabolomics, and other deep molecular phenotypes will inherently define groups of patients that are similar with respect to disease-relevant clinical outcomes. Note that “outcome” is here defined in a statistical sense, and also includes parameters such as sex, prevalent disease and current BMI. Recent examples of molecularly-defined subgroups include the identification of subtypes of lymphoma that severely impact survival [4], subtypes of various cancers identified in the ‘The Cancer Genome Atlas’ [5], and patient stratification in allergic diseases [6].

However, the identification of such subgroups in complex omics datasets with variety of patient measurements and clinical outcomes requires automated computational approaches. Adding to this complexity, patient subgroups might occur nested inside other, higher-level subgroups; for example, a group of high-risk patients within the male subpopulation or within a certain age group. The ‘SGI’ (subgroup identification) package implements a novel method to automatically generate omics-based subgroups of any granularity for an arbitrary number clinical variables of interest. It moreover provides a comprehensive set of methods to visualize the associations for further interpretation.

## 2 Description

### 2.1 SGI method

The SGI algorithm generates a hierarchical clustering of samples and runs a two-group association test against the analyzed clinical outcomes at each branching point in the tree. An example output plot is shown in Figure 1, which will be further discussed in the application section below. The algorithm works as follows: **(1) Clustering**. A dendrogram of the samples is generated using standard hierarchical clustering on the input data matrix. The function accepts any hclust object, giving the user full control over the choice of distance and linkage functions. (2) **Generate valid cluster pairs**. To avoid low-powered calculations in small clusters, the method enumerates all branching points where both left and right subclusters are above a user-defined size threshold. This results in a list of “valid” cluster pairs. SGI runs with a default setting of 5% of the sample size. (3) **Association analysis.** Run statistical tests with all clinical outcomes for all valid cluster pairs. SGI has built-in implementations for categorical outcomes (Fisher’s exact tests), continuous outcomes (two-sample t-tests) and survival outcomes (log-rank tests). The appropriate test is automatically determined by the toolbox based on the data type of the clinical variable. Furthermore, the user can define arbitrary association functions for more complex data types. **(4) Multiple testing correction.** Since a dendrogram clusters the samples into cascaded, non-overlapping groups, all statistical tests are strictly independent. Thus, SGI performs Bonferroni multiple testing correction by adjusting each p-value by a factor of the number of valid cluster pairs.

**Figure 1:**
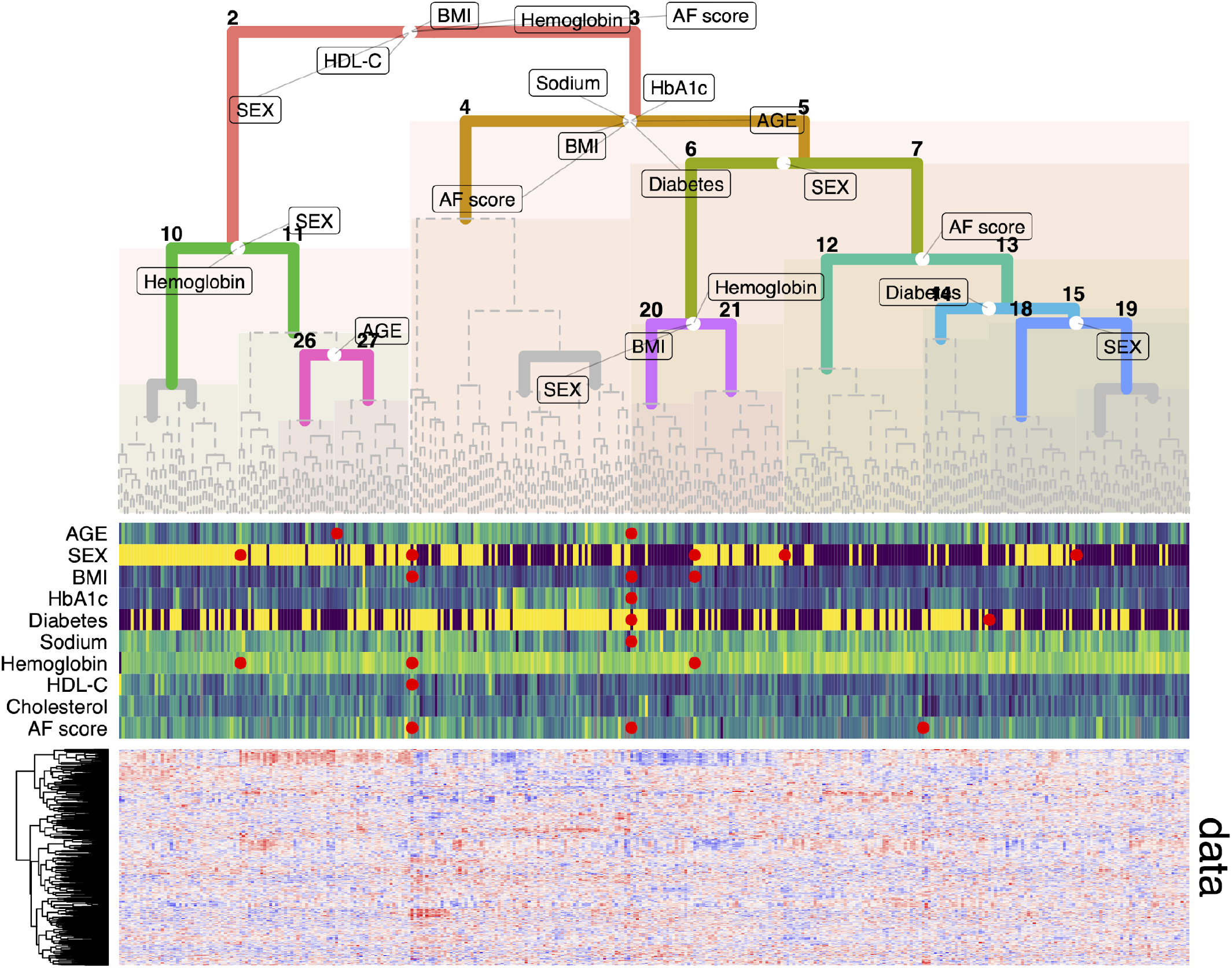
Application example. Blood metabolomics-based clustering of n=356 participants of the QMdiab study. White circles in the tree indicate the left/right splitting points of the samples in the data (note that these are not centered if the subclusters are of unequal size). Markings on the tree indicate statistically significant associations of the parameter with the respective left and right subgroups at that split. Heatmap track below the tree shows individual values for selected parameters. Red circles between gaps indicate significant results for left vs. right at that split and are horizontally aligned with their respective white circles on the tree. Bottom panel shows metabolomics data matrix behind the clustering. DIAB=diabetes diagnosis yes/no.

In addition to its core functionality, SGI provides support functions to extract, plot, print, and summarize all clustering and association results and test statistics, which allows the user to access all intermediate results to further analyze the subgroup results.

### 2.2 Visualization

Association results with multiple outcomes on a hierarchical tree are inherently complex to visualize. The SGI toolbox provides a variety of dedicated functions to visually inspect the statistical associations. This includes tree visualizations of all statistically significant outcome associations that are displayed at the respective branching points and heatmaps of the actual data (Figure 1). Moreover, the user can generate plots to inspect specific associations, e.g., boxplots of a quantitative clinical outcome between two clusters.

This allows the user to obtain a quick overview of the correlation structure between the input omics dataset, the resulting patient groups, and the clinical features that are analyzed. The simultaneous visualization of all data also allows to dissect the relationship between confounding variables, avoiding the predefined choice of a list of confounder variables to correct for.

### 2.3 Multi-omics datasets

The SGI package provides clustering functionality for the analysis of multi-omics datasets, i.e., datasets where more than one omics technology has been measured for the same samples. To this end, SGI generates a joint samples X sample distance matrix from the individual distance matrices of each omics layer. Since different omics layers will have varying numbers of variables, the respective distance values are not at comparable scales. The toolbox thus normalizes each individual distance matrix by its maximum and defines *D* = *D*_1_/*max* (*D*_1_) + *D*”/*max* (*D*”) + ⋯ + *D_l_*/*max* (*D_l_*), with *D* representing the final distance matrix, *D*_1_ etc. the original distance matrices and *l* the number of omics datasets. The approach was adapted from [7], where it was originally introduced to generate a Ward-like clustering. Notably, the method also works to normalize multi-omics contributions for other types of linkages, such as average linkage and complete linkage. Moreover, the method works with any distance metric. A detailed example of the multi-omics capabilities of SGI on combined plasma, urine, and saliva metabolomics data can be found in the example R codes of the github repository.

## 3 Application example

We demonstrate the functionality of the SGI package on plasma metabolomics data set from the ‘QMdiab’ diabetes case/control study with 356 participants [8,9]. Outcome parameters were type 2 diabetes diagnosis and nine anthropometric and clinical biochemistry parameters: age, sex, BMI, HbA1c, albumin, hemoglobin, LDL cholesterol, total cholesterol and skin auto-fluorescence [10]. The goal was to determine clusters of study participants defined by their profiles of circulating metabolites, and how these metabolomic clusters correlate with the different clinical parameters. The resulting visualization is shown in Figure 1. The following lines of code directly generate that plot using the SGI package:

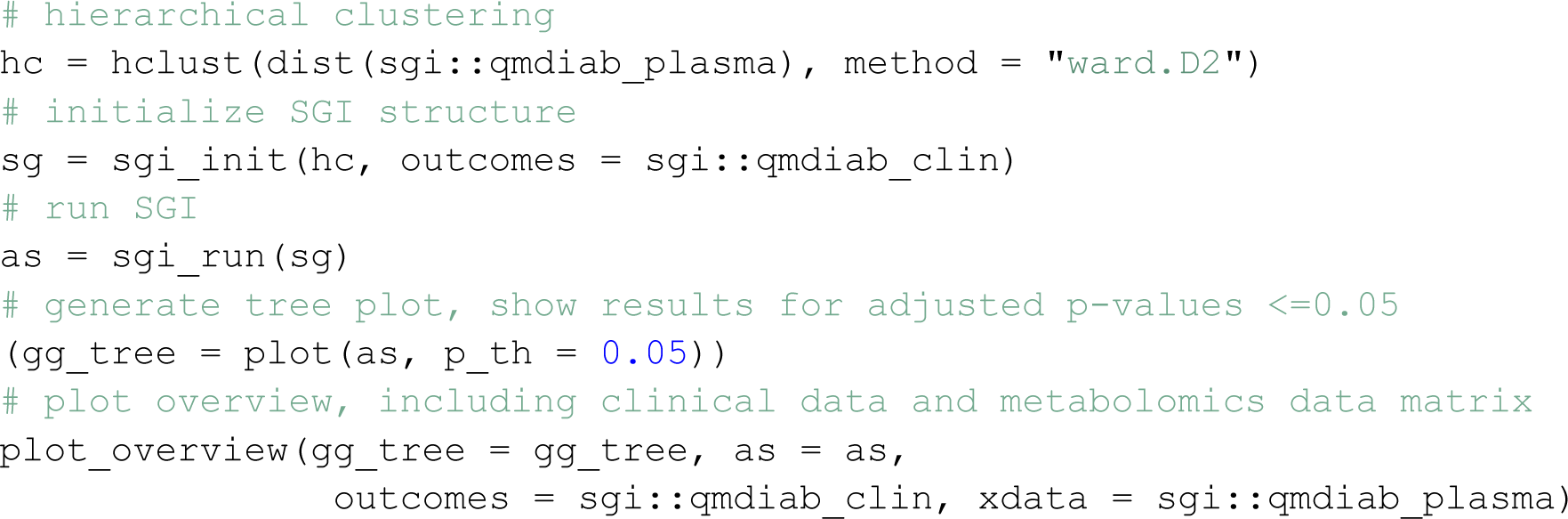

In this R code, `sgi::qmdiab_plasma`and `sgi::qmdiab_clin`are data frames holding the metabolomics and clinical variables, respectively. These data frames are contained in the package. The tree shows how metabolomic profiles separate study participants into two major groups at the top level of the tree (clusters 2 vs. 3), one with older individuals and a higher proportion of diabetes, and a younger group with lower prevalence of diabetes. Inside those two groups, further subgroups were identified, for example a male vs. female cluster which is also defined by significant differences of BMI inside the healthier group (clusters 6 vs. 7).

## 4 Conclusion

SGI provides an unbiased, data-driven way to automatically identify sample subgroups in omics profiles. It identifies and visualizes complex, hierarchical relationships for an arbitrary number of clinical outcomes in a visually intuitive way. The toolbox is easy to use, open source, and comes with a series of examples in the online repository.

## Acknowledgements & Funding

KS is supported by ‘Biomedical Research Program’ funds at Weill Cornell Medical College in Qatar, a program funded by the Qatar Foundation and multiple grants from the Qatar National Research Fund (QNRF). JK is supported by the National Institute of Aging of the National Institutes of Health under award 1U19AG063744.

## References

1. Rouzier R, Perou CM, Symmans WF, Ibrahim N, Cristofanilli M, Anderson K, et al. Breast cancer molecular subtypes respond differently to preoperative chemotherapy. Clin Cancer Res. 2005;11: 5678– 5685. doi:10.1158/1078-0432.CCR-04-2421

2. Eddy S, Mariani LH, Kretzler M. Integrated multi-omics approaches to improve classification of chronic kidney disease. Nature Reviews Nephrology. Nature Research; 2020. pp. 657–668. doi:10.1038/s41581-020-0286-5

3. Collisson EA, Sadanandam A, Olson P, Gibb WJ, Truitt M, Gu S, et al. Subtypes of pancreatic ductal adenocarcinoma and their differing responses to therapy. Nat Med. 2011;17: 500–503. doi:10.1038/nm.2344

4. Nowakowski GS, Czuczman MS. ABC, GCB, and Double-Hit Diffuse Large B-Cell Lymphoma: Does Subtype Make a Difference in Therapy Selection? Am Soc Clin Oncol Educ B. 2015; e449–e457. doi:10.14694/edbook_am.2015.35.e449

5. Sanchez-Vega F, Mina M, Armenia J, Chatila WK, Luna A, La KC, et al. Oncogenic Signaling Pathways in The Cancer Genome Atlas. Cell. 2018;173: 321-337.e10. doi:10.1016/j.cell.2018.03.035

6. Agache I, Akdis CA. Precision medicine and phenotypes, endotypes, genotypes, regiotypes, and theratypes of allergic diseases. Journal of Clinical Investigation. American Society for Clinical Investigation; 2019. pp. 1493–1503. doi:10.1172/JCI124611

7. Chavent M, Kuentz-Simonet V, Labenne A, Saracco J. ClustGeo: an R package for hierarchical clustering with spatial constraints. Comput Stat. 2018;33: 1799–1822. doi:10.1007/s00180-018-0791-1

8. Mook-Kanamori DO, El-Din Selim MM, Takiddin AH, Al-Homsi H, Al-Mahmoud KAS, Al-Obaidli A, et al. 1,5-Anhydroglucitol in saliva is a noninvasive marker of short-term glycemic control. J Clin Endocrinol Metab. 2014;99. doi:10.1210/jc.2013-3596

9. Do KT, Pietzner M, Rasp DJ, Friedrich N, Nauck M, Kocher T, et al. Phenotype-driven identification of modules in a hierarchical map of multifluid metabolic correlations. npj Syst Biol Appl. 2017;3: 28. doi:10.1038/s41540-017-0029-9

10. Mook-Kanamori MJ, El-Din Selim MM, Takiddin AH, Al-Homsi H, Al-Mahmoud KAS, Al-Obaidli A, et al. Ethnic and gender differences in advanced glycation end products measured by skin auto-fluorescence. Dermatoendocrinol. 2013;5: 325–330. doi:10.4161/derm.26046

